# BAP1 modulates endoplasmic reticulum stress signaling and balances liver homeostasis and malignant progression

**DOI:** 10.1101/2025.11.06.686752

**Authors:** Olatunji Oluwole Gege, Agnieszka Seretny, Luise Butthof, Long Pan, Lena Wendler-Link, Leonardo Traini, Lio Boese, Sabrina Neuhaeuser, Alexander Scheiter, Danijela Heide, Christian Luechtenborg, Britta Bruegger, Diego Calvisi, Mathias Heikenwaelder, Jessica Zucman-Rossi, Marco Breining, Darjus Felix Tschaharganeh

## Abstract

**Objective:** Primary liver cancer is a leading cause of cancer-related mortality and harbors recurrent mutations in chromatin regulators such as BRCA1-associated protein 1 (BAP1), yet their functional impact remains unclear. We investigated how BAP1 deficiency affects liver homeostasis and tumorigenesis to clarify its functional role.

**Design:** We employed inducible, liver-specific BAP1 knockdown in mice subjected to diet-induced metabolic stress (including rescue experiments), alongside autochthonous hydrodynamic CRISPR models, and profiled livers by RNA-seq, immunohistochemistry, and mass spectrometry-based lipidomics. Complementary mechanistic assays in liver cancer cells examined the unfolded protein response (UPR) under endoplasmic reticulum (ER) stress; findings were supported by immunohistochemical and transcriptomic analyses of BAP1-mutant patient samples.

**Results:** BAP1 safeguards liver homeostasis under diet-induced metabolic stress, as its loss triggers ER stress, hepatocyte death, and acute liver failure. Lipidomics revealed a shift toward ER-stress-associated dyslipidemia, and transcriptomics showed negative enrichment of fatty-acid metabolism and positive enrichment of UPR pathways. In contrast, BAP1 loss synergizes with oncogenic drivers to accelerate tumorigenesis in autochthonous liver cancer models, underscoring its context-dependent tumor suppressor function. Mechanistically, BAP1 directly regulates the ER stress mediator *DDIT3* (CHOP) through chromatin remodeling, linking BAP1 loss to maladaptive stress responses. Consistently, elevated *CHOP* expression was observed in BAP1-mutant human liver cancers and other tumor types.

**Conclusion:** These findings establish BAP1 as a key chromatin regulator that connects stress adaptation to both liver homeostasis and tumorigenesis, highlighting the BAP1-UPR axis for future translational assessment.

**What is already known on this topic?:** BAP1 is a recurrently mutated chromatin regulator across cancers, including primary liver cancer, but its functional role in liver biology and tumorigenesis has remained unclear. Metabolic dysfunction-associated liver disease is a growing driver of liver tumorigenesis and is characterized by lipid imbalance, ER stress, and activation of the unfolded protein response.

**What this study adds?:** We show that BAP1 is a key chromatin regulator that integrates metabolic and ER stress responses in the liver. Loss of BAP1 undermines cellular adaptation to metabolic challenge and cooperates with oncogenic signals to promote liver tumorigenesis via dysregulated *DDIT3* (CHOP) expression, linking chromatin control to hepatic stress resilience and disease progression.

**How this study might affect research, practice or policy?:** Our findings position the UPR-CHOP axis as a candidate therapeutic vulnerability in BAP1-deficient liver cancers, particularly in the context of MASLD/MASH, and provide a conceptual framework for targeting stress adaptation pathways in precision oncology.

## Introduction

Primary liver cancer remains a leading cause of cancer mortality, driven by late detection, limited therapeutic options, and pronounced genetic heterogeneity [1,2]. Recent sequencing efforts have uncovered a complex mutational landscape in liver cancer, with frequent alterations in *TERT, TP53*, and *CTNNB1*, as well as recurrent mutations in chromatin modifiers such as *ARID1A, PBRM1*, and *BAP1* [3,4]. Despite their prevalence, the functional relevance of many of these mutations in liver tumorigenesis remains poorly understood.

BRCA1-associated protein 1 (BAP1) encodes a deubiquitinase that regulates transcription, chromatin architecture [5,6], and DNA repair [7]. BAP1 removes monoubiquitin from H2AK119Ub, antagonizing Polycomb-mediated repression and facilitating gene activation through the PR-DUB complex [8]. It also stabilizes the BRCA1/BARD1 complex, helping maintain genome integrity⍰[9]. Beyond these roles, BAP1 exerts context-dependent effects on cell death pathways, including both apoptotic protection and ferroptotic priming⍰[10,11].

Somatic *BAP1* mutations occur across diverse malignancies, including uveal melanoma, mesothelioma, and renal cell carcinoma⍰[8,12]. Interestingly, the prognostic significance of BAP1 loss varies by tissue; it predicts improved outcomes in mesothelioma [13] but poor survival in renal cancer⍰[14]. In liver cancer, frequent inactivation of BAP1 has been documented, yet its mechanistic role remains unclear.

This gap is particularly important in light of the growing global burden of metabolic dysfunction-associated steatotic liver disease (MASLD) and its progressive form, metabolic dysfunction-associated steatohepatitis (MASH), now recognized as major drivers of hepatocellular carcinoma [15,16]. While MASLD and MASH are marked by lipid overload, ER stress, and persistent unfolded protein response (UPR) activation [17,18], how metabolic stress interfaces with tumor-relevant genetic mutation remains poorly understood. Chromatin regulators like BAP1 are well positioned to integrate these signals, yet their role in stress adaptation remains largely unexplored.

Here, we employed complementary liver-specific mouse models to investigate the role of BAP1 in liver biology, pathology, and tumorigenesis. Using diet-induced metabolic stress models that recapitulate key features of MASLD/MASH [19,20], we demonstrate that *BAP1* loss sensitizes the liver to metabolic stress, disrupts lipid homeostasis, activates ER stress pathways, and ultimately leads to acute liver damage and death. In contrast, under tumor-promoting conditions BAP1 depletion accelerates liver cancer development. In the oncogenic setting, unresolved ER stress in BAP1-deficient livers appears to shift from a pro-death signal to a driver of tumor progression. These findings position BAP1 as a critical regulator of hepatic stress resilience and highlight its role in tumor suppression.

## Results

### A novel inducible, liver-specific model for BAP1 suppression

To investigate the role of BAP1 in liver biology, we used a transgenic mouse model that enables inducible, liver-specific knockdown of genes of interest [21,22]. This system employs potent short hairpin RNAs (shRNAs) under the control of tetracycline-responsive elements (TRE) enabling temporal control of gene silencing with doxycycline (Dox) [23]. A lox-Stop-lox cassette preceding the reverse-tetracycline-transactivator (rtTA3) allows for spatially restricted activation of the TRE promoter when crossed with a tissue-specific Cre recombinase line. For liver specificity, we used Albumin-Cre [24]. The model includes two reporters: mKate2 marks cells with Cre-mediated recombination, and GFP indicates shRNA induction upon Dox treatment, providing clear readouts of both events. (**Figure 1**).

**Figure 1.**
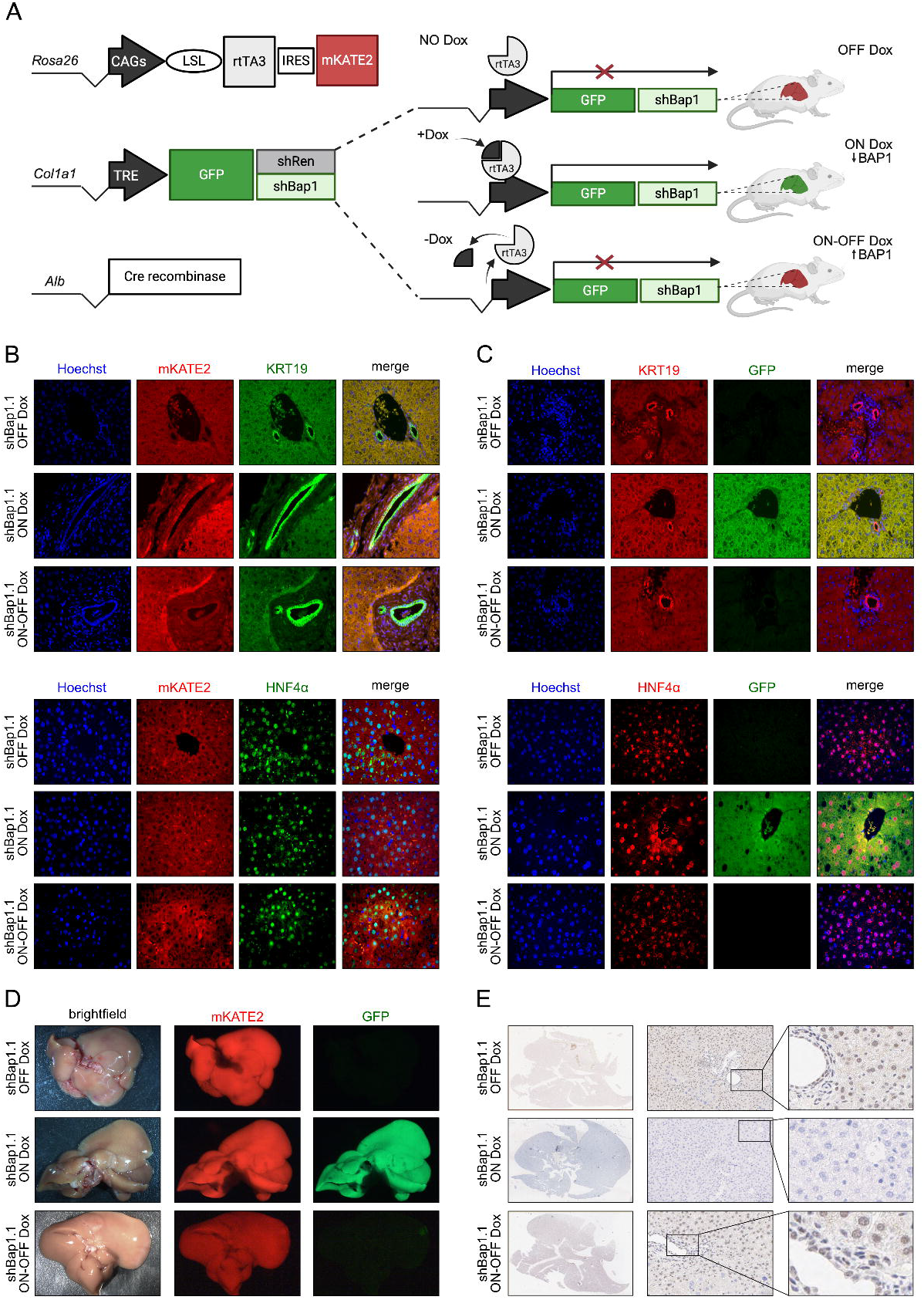
Development and validation of the doxycycline-inducible liver-specific shBap1 mouse model. (A) Schematic of the transgenic system combining Cre-dependent rtTA3/mKate2 expression at the *Rosa26* locus with TRE-driven GFP-shRNA integration at the *Col1a1* locus. (B&C) Representative immunofluorescence images at the 6-week time point (40x). (B) Co-localization of mKate2 (red) in KRT19^+^ (green) cholangiocytes (upper panel) and HNF4α^+^ (green) hepatocytes (lower panel), indicating targeting of both liver epithelial cell types. (C) Co-localization of GFP (green) with KRT19 (red) and HNF4α (red) confirming shRNA induction in both cholangiocytes and hepatocytes. (D) Representative images of a dissected liver showing mKate2 (red), marking Cre recombination, and GFP (green), indicating doxycycline-induced shRNA expression. (E) Representative immunohistochemical images for BAP1 confirm loss of nuclear BAP1 in ON Dox livers; panels show whole-slide overview (left), 10× field (middle), and 40× field (right).

We generated two distinct shBAP1 mouse strains, each harboring a single potent shRNAs targeting BAP1 (shBAP1.1 and shBAP1.2), as well as a complementary shRenilla mouse strain targeting Renilla luciferase as a negative control (**Figure 1A**). Using three doxycycline regimens: ON (continuous Dox), OFF (no Dox), and ON-OFF (2 weeks on Dox, followed by 4 weeks off Dox), we assessed inducibility and reversibility in major liver epithelial cell types. Immunofluorescence confirmed mKate2 expression in both cholangiocytes (KRT19^+^) and hepatocytes (HNF4α^+^), demonstrating successful targeting of both cell types (**Figure 1B**). To examine the spatial distribution of shRNA expression, we co-stained for GFP with KRT19 or HNF4α (**Figure 1C**). These analyses confirmed shRNA induction in both cholangiocytes and hepatocytes, with GFP signal co-localizing with KRT19^+^ and HNF4α^+^ cells.

Across these regimens, robust mKate2 was present in all groups, consistent with Cre-mediated recombination, while GFP fluorescence was restricted to Dox-treated animals (and declined after withdrawal), validating tight control of shRNA induction (**Figure 1D**). Immunohistochemistry for BAP1 confirmed Dox-dependent loss of nuclear signal (**Figure 1E**). Together, these data validate the system as a robust and tightly regulated tool for liver-specific modulation of BAP1 expression.

To assess long-term knockdown efficiency, we examined livers from Dox-treated shBap1 mice at 6 and 24 weeks. Western blot analysis confirmed stable BAP1 protein depletion over time, with GFP expression correlating with shRNA induction (**Supplementary Figure 1A**). Notably, BAP1 knockdown was also evident at the transcript level (**Supplementary Figure 1B**). Serum ALT (alanine aminotransferase), AST (aspartate aminotransferase), and bilirubin levels, established indicators of liver injury [25–27], remained stable across all groups and timepoints (**Supplementary Figure 2A-C**), indicating preserved liver function. Histological analysis showed largely preserved liver architecture (**Supplementary Figure 2D**).

Overall, these findings indicate that prolonged BAP1 depletion in the liver causes only mild histological changes without impairing liver function or promoting tumorigenesis under basal conditions.

### BAP1 determines liver resilience and survival under metabolic stress

To assess BAP1 function in metabolic liver injury and subsequent tumor development, we used a dietary-induced Metabolic Dysfunction-Associated Steatohepatitis (MASH) approach, challenging mice with a choline-deficient, high-fat diet (CD-HFD) under ON- or OFF-Dox conditions [15,23,28]. This diet imposes a well-characterized metabolic burden on the liver, modeling nutrient imbalance and excess, particularly in fat intake, that contribute to steatohepatitis and liver cancer in humans [29,30]. Surprisingly, Dox-treated shBap1 mice displayed rapid onset of lethality, with over 70% of shBap1.1 and 60% of shBap1.2 mice reaching humane endpoints by week 6, whereas all Dox-treated shRenilla control mice survived (**Figure 2A**).

**Figure 2.**
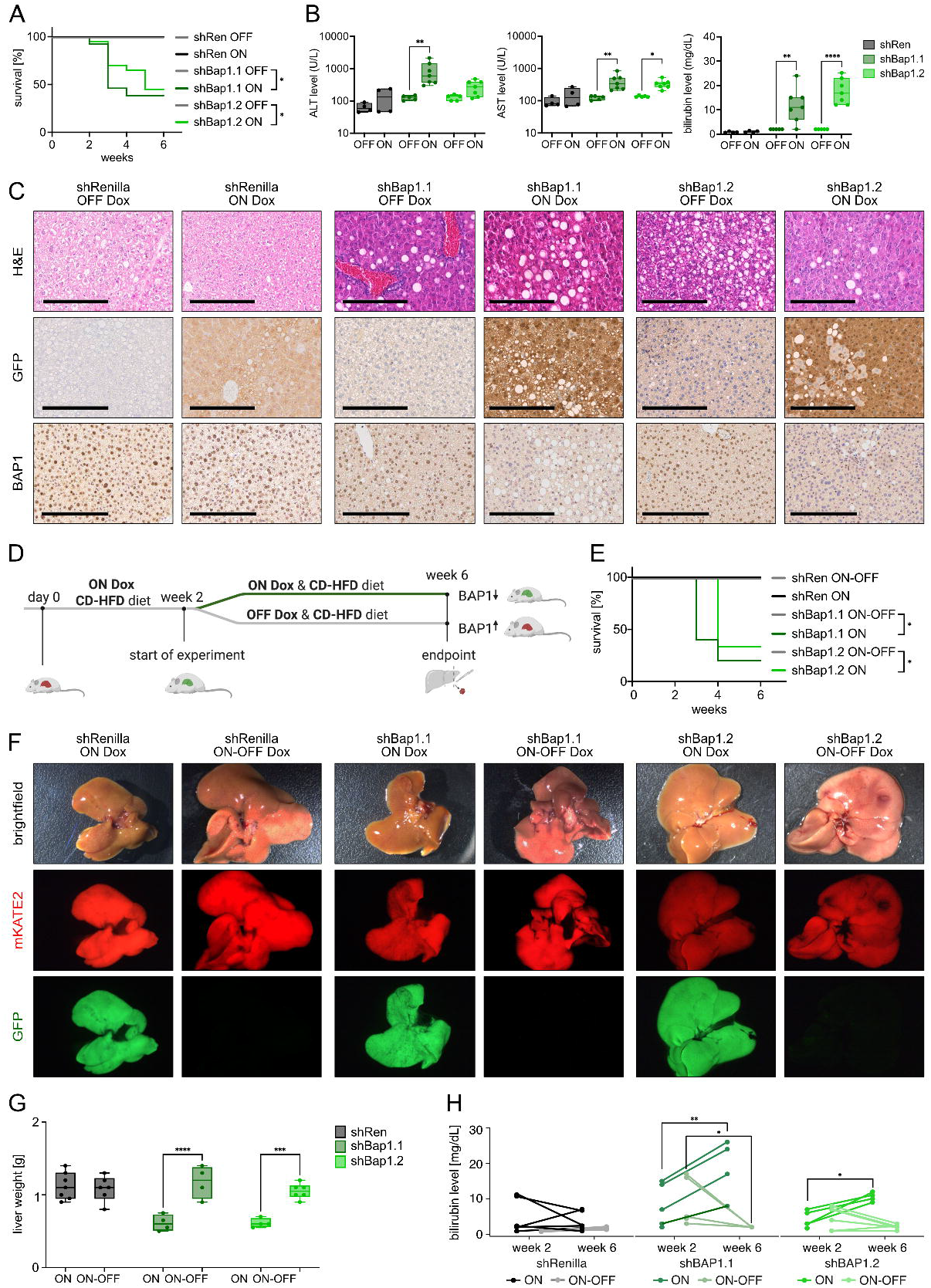
BAP1 loss precipitates acute liver failure under metabolic stress, reversible upon restoration. (A) Kaplan-Meier survival curves of shBap1.1, shBap1.2, and shRenilla mice fed CD-HFD under continuous Dox (ON) or no Dox (OFF). N ≥ 7, Log-rank (Mantel-Cox) test. (B) Endpoint serum levels of ALT, AST, and bilirubin show elevated liver injury markers in BAP1-deficient mice. N ≥ 4, two-way ANOVA with Šídák’s test. (C) Representative liver sections stained with H&E (top), GFP (middle), and BAP1 (bottom). Dox-treated shBap1 livers exhibit steatosis, robust GFP signal, and loss of nuclear BAP1, scale bar: 200 µm. (D) Experimental design of Dox withdrawal rescue (ON-OFF) with continuous CD-HFD throughout the entire 6-week period. (E) Kaplan-Meier survival curves showing improved survival upon Dox withdrawal in BAP1-deficient mice, indicating reversibility. N ≥ 5, Log-rank (Mantel-Cox) test. (F) Representative brightfield and fluorescence liver images at endpoint. Reduced GFP signal in ON-OFF groups indicates transgene silencing and restoration of BAP1 expression. (G) Quantification of liver weight at endpoint. ON-OFF mice maintain liver mass compared to ON groups, consistent with recovery. N ≥ 4, two-way ANOVA with Šídák’s test. (H) Serum bilirubin levels at week 2 and week 6 in ON and ON-OFF mice. Decline upon Dox withdrawal reflects improved liver function. N ≥ 4, two-way ANOVA with Šídák’s test.

Serum analyses revealed marked increase in ALT, AST, and bilirubin in BAP1-deficient mice, consistent with acute liver damage and impaired function as a likely cause of lethality. Values remained within normal ranges in shRenilla controls regardless of Dox exposure (**Figure 2B**). Histological and immunohistochemical analyses revealed liver injury in BAP1-deficient mice, characterized by hypereosinophilic, apoptotic hepatocytes and loss of tissue integrity in hematoxylin-eosin (H&E) staining (**Figure 2C**). GFP staining verified robust shRNA induction in all Dox-treated groups, while BAP1 immunostaining revealed efficient nuclear depletion in ON Dox shBap1.1 and shBap1.2 livers. In contrast, shRenilla controls maintained BAP1 expression and preserved hepatic architecture with no overt pathology, confirming a specific effect of BAP1 loss under metabolic stress.

To disentangle the contribution of choline deficiency on the observed phenotype, we subjected mice to a high-fat diet (HFD) without choline restriction. This regimen models features of Metabolic Dysfunction-Associated Steatotic Liver Disease (MASLD), with inflammatory changes that are generally milder than those induced by CD-HFD conditions [31–33]. Even under these less severe conditions, more than 70% of shBap1.2 and over 30% of shBap1.1 mice became moribund, while all shRenilla controls survived (**Supplementary Figure 3A**). Dox-treated shBap1 mice exhibited elevated AST and bilirubin levels, indicating liver injury and dysfunction compared to shRenilla mice (**Supplementary Figure 3B**).

Finally, we leveraged the spatiotemporal control provided by our mouse model to assess the reversibility of the phenotype following BAP1 restoration. We first induced BAP1 deficiency by Dox administration in animals on CD-HFD. After two weeks, animals were divided into two cohorts: one continued on Dox (Bap1-deficient) and the other had Dox withdrawn (Bap1-restored) (**Figure 2D**). This intervention markedly improved survival in both shBap1.1 and shBap1.2 mice (**Figure 2E**). Reduced GFP signal confirmed transgene silencing and BAP1 restoration (**Figure 2F**). Consistent with functional recovery, liver weight was preserved compared with mice with sustained BAP1 loss (**Figure 2G**), and serum bilirubin levels normalized, indicating restored liver function (**Figure 2H**).

Together, these findings show that BAP1 loss accelerates liver damage in response to metabolic stress, underscoring its critical role in maintaining liver integrity and homeostasis.

### BAP1 loss triggers ER stress and lipid imbalance under nutrient overload

To investigate the mechanisms underlying liver damage and fatality in BAP1-deficient mice under metabolic stress, we first explored whether immune cell infiltration or inflammatory cell death contributed to the rapid deterioration.

Immunohistochemistry for F4/80 (macrophage marker [34]), CD3 (T cells marker [35]), and RIPK3 (necroptosis marker [36]) showed no major differences between groups (**Supplementary Figure 4A&B**), suggesting that neither immune recruitment nor necroptotic signaling played a significant role. In contrast, TUNEL staining [37] revealed increased DNA fragmentation in Dox-treated shBap1.1 and shBap1.2 livers, but not in controls (**Figure 3A; Supplementary Figure 4C**). This coincided with nuclear BAP1 loss, supporting a link between BAP1 deficiency and apoptotic or genotoxic stress.

**Figure 3.**
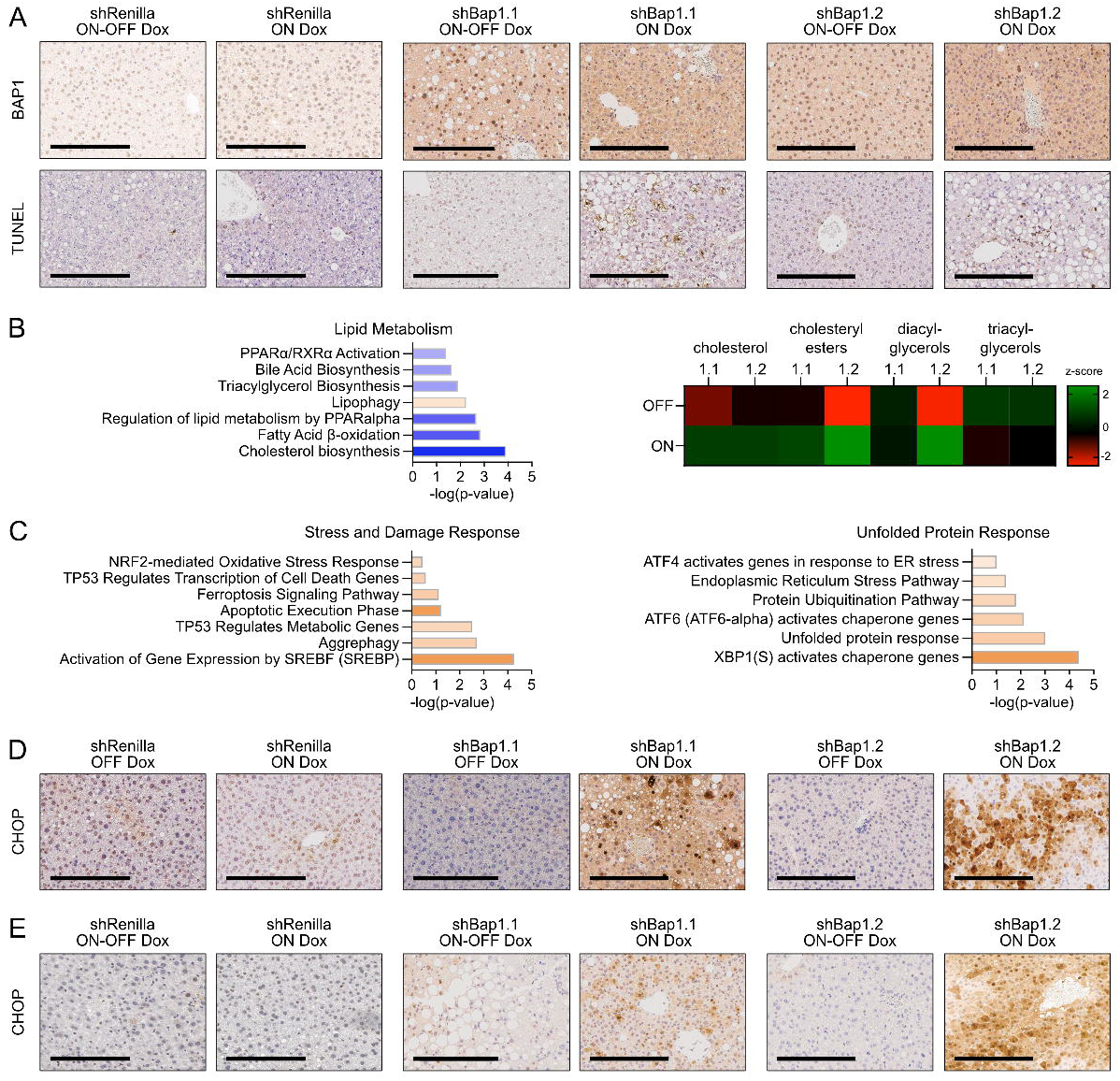
BAP1 loss induces ER stress and lipid imbalance under dietary challenge. (A) Representative immunohistochemistry images for BAP1 and TUNEL show nuclear BAP1 loss in Dox-treated shBap1 livers and increased apoptosis; scale bar: 200⍰lµm. (B) RNAseq (left) and lipidomics (right) reveal altered lipid metabolism in BAP1-deficient livers under CD-HFD, N = 4-5. Lipidomics presented as z-score mean values. (C) Top differentially enriched pathways from Ingenuity Pathway Analysis in CD-HFD-fed BAP1-deficient livers highlight ER stress UPR. (D&E) Representative immunohistochemical images of CHOP staining demonstrate (D) strong ER stress activation in shBap1 livers under Dox; (E) CHOP expression is reversed in ON-OFF rescue mice, confirming restoration of BAP1 alleviates ER stress. Scale bar: 200⍰lµm.

To further investigate the underlying mechanisms of liver cell death, we performed RNA sequencing on BAP1-deficient livers exposed to CD-HFD. Transcriptomic analysis revealed impaired lipid metabolism, activation of oxidative and ER stress responses, and induction of TP53-mediated apoptosis (**Figure 3B&C, Supplementary Figure 5A**). These data suggest that BAP1 loss compromises hepatic stress resilience primarily through intrinsic pathways rather than inflammation or immune-mediated injury.

Lipidomics analysis of CD-HFD-fed shBap1.1 and shBap1.2 mice corroborated the transcriptomic findings, revealing disrupted hepatic lipid homeostasis (**Figure 3B**). BAP1 deficiency causes a metabolic bottleneck: decreased degradation and export of lipids, with increased lipophagy most likely attempting to mitigate accumulation, culminating in lipid buildup and metabolic stress. These data collectively indicate that BAP1 loss disrupts lipid metabolism at both the transcriptional and metabolic levels under dietary stress.

Interestingly, we observed strong upregulation of the unfolded protein response (UPR) alongside the marked downregulation lipid metabolism in Dox-treated shBap1 mice (**Figure 3C**). Notably, ER stress markers were among the most enriched transcripts, whereas key lipid metabolism genes were significantly repressed in BAP1-deficient group.

Given the established connection between lipid dysregulation and UPR activation [38,39], along with our transcriptomic evidence for both, we next assessed the severity of ER stress. We focused on CHOP, encoded by *DDIT3*, a key pro-apoptotic UPR effector [40], which ranked among the top upregulated transcripts in BAP1-deficient livers (**Supplementary Figure 5B**). CHOP immunohistochemistry revealed strong staining in livers of shBap1.1 and shBap1.2 mice, while shRenilla controls showed minimal induction (**Figure 3D**). Importantly, CHOP expression was reversed in rescue experiments upon BAP1 re-expression, confirming the functional link between BAP1 loss and ER stress activation (**Figure 3E**).

Together, these findings demonstrate that BAP1 loss under metabolic stress compromises hepatic stress resilience by disrupting lipid metabolism and activating ER stress pathways, with CHOP serving as a key effector.

### BAP1 functions as a context-dependent tumor suppressor in the liver

While the combination of BAP1 loss with diet-induced metabolic stress caused acute liver failure in our experiments, each factor alone has been linked to liver tumorigenesis [30]. This prompted us to investigate whether BAP1 loss promotes liver tumors in a context-dependent manner, particularly when combined with other oncogenic alterations.

To test this, we used hydrodynamic tail vein injection (HTVI) to deliver CRISPR/Cas9 constructs directly into the liver [41,42], targeting *Bap1* in combination with known liver cancer oncogenes or tumor suppressors.

First, we assessed whether BAP1 inactivation cooperates with YAP activation, a well-established oncogenic effector of the Hippo pathway frequently dysregulated in HCC [43]. Co-expression of constitutively active mutant YAP (YAP^S127A^) with sgBap1 led to early mortality and aggressive tumor formation (**Figure 4A**). Four of five YAP;sgBap1 mice developed multifocal liver tumors within 16 weeks, and both liver weight and liver-to-body weight ratios were significantly increased, highlighting enhanced tumor burden in the BAP1-deficient setting. These data demonstrate that BAP1 loss accelerates YAP-driven liver tumorigenesis.

**Figure 4.**
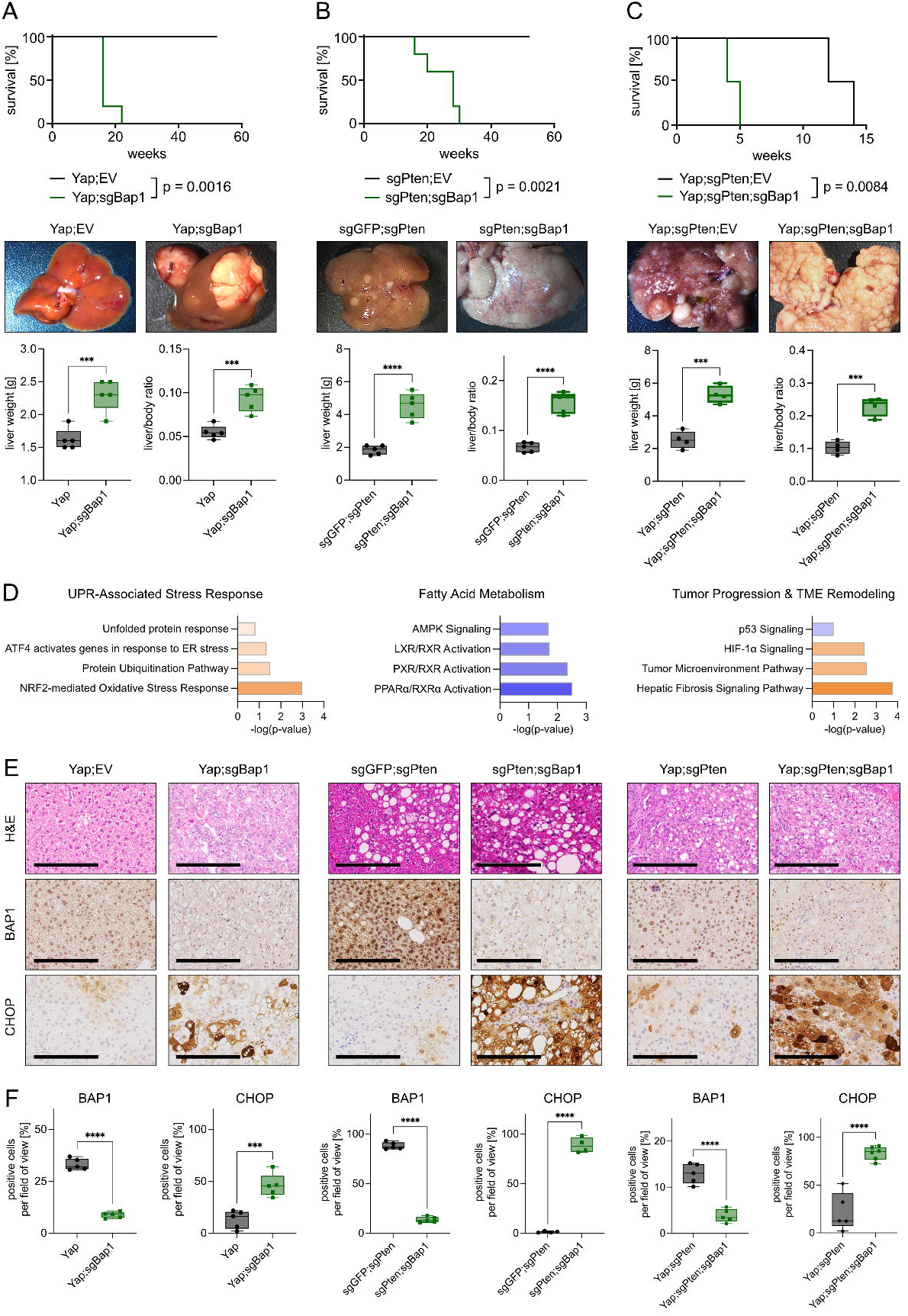
BAP1 loss accelerates liver tumorigenesis and induces ER stress in oncogenic models. (A-C) BAP1 deficiency shortens survival and increases liver tumor burden across different oncogenic contexts. Shown are Kaplan-Meier survival curves (top), representative liver images (middle), and liver-to-body weight ratios (bottom) from hydrodynamic tail vein injection models: (A) YAP;EV and YAP;sgBap1, (B) sgGFP;sgPten and sgPten;sgBap1, and (C) YAP;sgPten and YAP;sgPten;sgBap1. N = 5, survival: log-rank (Mantel-Cox) test; liver weight comparisons: unpaired two-tailed Student’s t-test. (D) Top enriched pathways from transcriptomic analysis of BAP1-deficient tumors indicate activation of the UPR, repression of fatty acid metabolism, and upregulation of tumor-promoting signaling pathways. (E) Representative histological and immunohistochemical images (H&E, BAP1, CHOP) show altered morphology, nuclear BAP1 loss, and strong CHOP induction in BAP1-deficient tumors. Scale bar: 2001lµm. (F) Quantification of BAP1^+^ and CHOP^+^ cells. N = 5, unpaired two-tailed Student’s t-test.

We next tested whether BAP1 inactivation cooperates with loss of tumor suppressors. PTEN, a key negative regulator of PI3K-AKT signaling, is frequently lost in liver cancer [44–46]. Mice injected with sgPten and sgBap1 developed hepatomegaly and became moribund about 30 weeks post-injection, while sgPten-only controls remained asymptomatic, with only occasional small nodules observed at endpoint (**Figure 4B**). Liver weights and liver-to-body weight ratios were significantly elevated in the double-knockout cohort, indicating synergy between BAP1 and PTEN loss.

Given that PTEN loss activates YAP signaling in the liver [47], we next examined whether BAP1 deficiency exacerbates tumorigenesis when both PI3K and Hippo pathways are simultaneously perturbed. Triple perturbation with YAP^S127A^, sgPten, and sgBap1 markedly accelerated tumor onset compared with the YAP^S127A^;sgPten control group (**Figure 4C**). Livers from the triple-targeted mice were enlarged and densely populated with tumor nodules, whereas control mice exhibited fewer lesions. Again, liver weight and liver-to-body weight ratios were significantly increased in the BAP1-deficient group.

Together, these data identify BAP1 as capable of cooperating with either YAP activation or PTEN loss, and further intensifying tumorigenesis when both pathways are engaged.

To characterize the molecular changes accompanying this transformation, we performed transcriptomic analysis of sgPten;sgBap1 tumors. Pathway enrichment revealed activation of UPR, oxidative stress responses, and tumor microenvironment remodeling, together with downregulation of fatty acid metabolism and PPAR signaling (**Figure 4D**). These alterations mirror those observed in BAP1-deficient livers under dietary challenge, suggesting a shared vulnerability axis driven by ER stress.

Histologically, BAP1-deficient tumors displayed HCC-like morphology and consistent nuclear loss of BAP1 protein (**Figure 4E**). Strikingly, CHOP was robustly upregulated in all BAP1-deficient tumors; strong CHOP staining was detected in YAP;sgBap1, sgPten;sgBap1, and YAP;sgPten;sgBap1 livers, whereas BAP1-proficient controls showed only minimal expression. These findings identify CHOP induction as a hallmark of BAP1 loss, extending its impact from metabolic dysfunction to malignant transformation.

### BAP1 deficiency drives CHOP upregulation and ER stress in human liver tumors

To determine whether the link between BAP1 loss and ER stress observed in mouse models extends to human liver cancer, we analyzed multiple liver cancer cell lines and patient-derived datasets. To phenocopy ER stress *in vitro*, we treated three independent cell lines (Huh7, Hep3B, and HepG2) with tunicamycin, a potent UPR inducer [48]. Under these conditions, BAP1 loss consistently led to elevated CHOP protein levels, indicating heightened sensitivity to ER stress (**Figure 5A-C**). Correspondingly, *DDIT3* transcript levels were significantly upregulated in BAP1-deficient cells upon tunicamycin exposure (**Figure 5D**).

**Figure 5.**
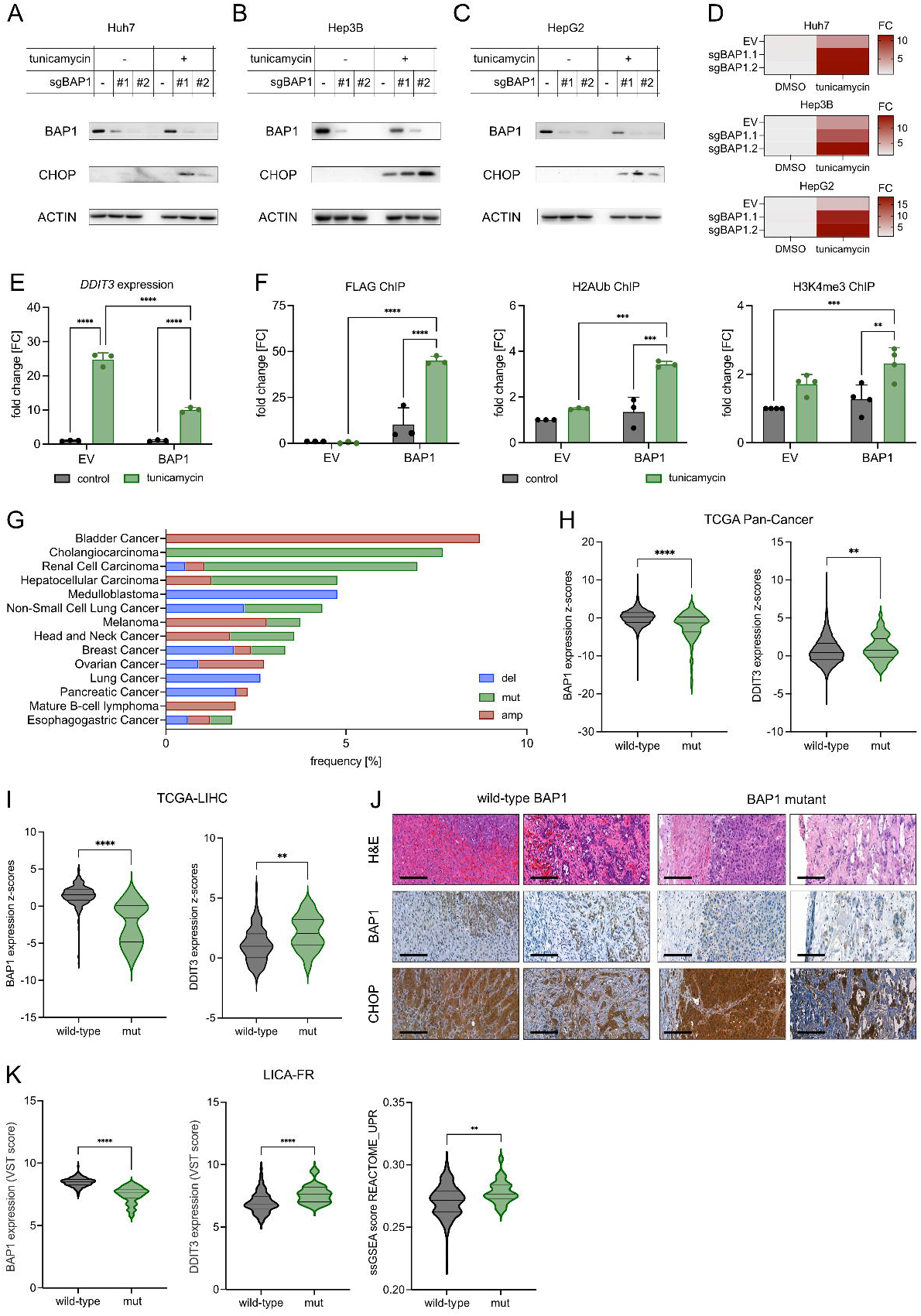
BAP1 loss triggers CHOP-driven unfolded protein response in human liver cancer models and patient samples. (A-C) Representative Western blot analysis showing elevated CHOP protein levels following tunicamycin-induced ER stress (10 µg/ml, 8-hour treatment) in BAP1 knockout HCC cell lines: (A) Huh7, (B) Hep3B, and (C) HepG2. N = 3. (D) Relative expression of *Ddit3* (CHOP) increases upon 24-hour tunicamycin (10 µg/ml) treatment in BAP1-deficient HCC cell lines (Huh7, Hep3B, HepG2) presented as fold change. N = 3, represented as fold change (FC). (E) Quantification of DDIT3 transcript levels in empty vector (EV) or BAP1-overexpressing cells with or without tunicamycin. One-way ANOVA with Tukey’s multiple comparisons test. (F) ChIP-qPCR analysis showing BAP1 binding at the *DDIT3* locus (FLAG) and corresponding changes in histone marks (H2Aub and H3K4me3) in EV and BAP1-overexpressing cells following tunicamycin treatment. One-way ANOVA with Tukey’s multiple comparisons test. (G) Frequency of BAP1 alterations: mutations (mut), deletions (del), and amplifications (amp) across cancer types in TCGA datasets. (H-I) Z-score analysis from publicly available patient datasets showing reduced *BAP1* and elevated *DDIT3* expression in BAP1-mutant tumors across all cancers (H; wild-type: N = 6552, mutant: N = 172) and specifically in liver cancer (I; wild-type: N = 328, mutant: N = 20). Violin plots (thick median line; quartiles shown), unpaired two-tailed Mann-Whitney’s U test. (J) Representative H&E and IHC images showing nuclear BAP1 loss and strong CHOP induction in BAP1-mutant human liver tumors (wild-type: N = 8, mut: N = 7). Scale bars: 200⍰µm. (K) ssGSEA analysis of human HCC RNA-seq showing decreased *BAP1* and elevated *DDIT3* expression, as well as increased unfolded protein response activity in BAP1-mutant (N = 27) compared to BAP1-wild-type (N = 379) tumors. VST score stands for variance stabilizing transform. Unpaired two-tailed Mann-Whitney’s U test.

To test whether BAP1 overexpression could dampen this response, BAP1 was overexpressed in HEK293T cells followed by tunicamycin treatment. This resulted in markedly lower *DDIT3* transcript levels compared with empty vector controls (**Figure 5E**), indicating a repressive role for BAP1 in CHOP regulation. Chromatin immunoprecipitation further demonstrated that BAP1 binds to the *DDIT3* locus under ER stress conditions. Intriguingly, this binding was accompanied by increased levels of both H2AUb and H3K4me3 (**Figure 5F**), suggesting establishment of a bivalent chromatin state [49–51]. The co-occurrence of repressive H2AUb and activating H3K4me3 may reflect a poised configuration, enabling stress-responsive regulation.

To evaluate the clinical relevance, we analyzed TCGA datasets [52,53]. BAP1 alterations were detected across multiple tumor types, with particularly high mutation frequency in cholangiocarcinoma, renal cell carcinoma, and hepatocellular carcinoma (**Figure 5G**). Notably, tumors harboring BAP1 mutations showed significantly lower *BAP1* and higher *DDIT3* expression levels compared with wild-type counterparts, both in pan-cancer datasets (**Figure 5H**) and within liver cancer (**Figure 5I**).

Furthermore, histological analysis of human liver tumors revealed an inverse relationship between nuclear BAP1 and CHOP levels. Tumors harboring BAP1 mutations showed widespread CHOP upregulation and loss of nuclear BAP1, whereas BAP1 wild-type tumors retained nuclear BAP1 and exhibited more restricted CHOP staining (**Figure 5J**). Finally, independent transcriptomic analysis of human HCCs (LICA-FR) (Pan *et al*., in submission) confirmed that BAP1-mutant tumors display features of heightened ER stress, with increased unfolded protein response activity and elevated DDIT3 expression, supporting that this relationship is evident at the transcriptomic level (**Figure 5K**).

Collectively, these findings establish BAP1 as a critical suppressor of ER stress in liver cancer, with its loss driving consistent CHOP upregulation across human models and patient tumors.

## Discussion

Primary liver cancer is a leading cause of cancer-related mortality worldwide, yet effective therapies remain limited [2]. This challenge is growing in urgency as metabolic dysfunction-associated steatotic liver disease (MASLD) and its progressive form, metabolic dysfunction-associated steatohepatitis (MASH), increasingly contribute to the global burden of hepatocellular carcinoma [30,54]. A deeper understanding of how genetic alterations influence tumor development and progression is essential to developing better treatment strategies. Yet, despite the fact that recurrent alterations in chromatin modifiers, such as BAP1, are frequent in hepatocellular carcinoma and intrahepatic cholangiocarcinoma [4,12], their functional impact remains poorly understood.

Here, we demonstrate that BAP1 safeguards hepatic resilience under metabolic stress and functions as a *bona fide* tumor suppressor in the liver. Loss of BAP1 sensitized hepatocytes to metabolic challenges, linking it to lipid imbalance, unresolved ER stress, and induction of the maladaptive transcription factor CHOP. Importantly, BAP1 restoration reversed stress signatures and improved survival, underscoring its protective role in maintaining hepatic homeostasis.

Dai *et al*. [55] showed *in vitro* that BAP1 deficiency elevates CHOP and that exogenous wild-type BAP1, under nutrient deprivation, is recruited to the *DDIT3* promoter together with a paradoxical increase in BAP1’s target mark, H2AK119Ub1. In our system, tunicamycin-induced UPR likewise resulted in BAP1 recruitment and elevated levels of H2AK119Ub1 at the *DDIT3* promoter, but we additionally observed elevated H3K4me3. Functionally, BAP1 overexpression reduced CHOP output upon UPR induction. Taken together, these data support a repressed-yet-poised (bivalent-like) promoter model [49–51], in which BAP1 occupancy appears to coincide with increased H2AK119Ub1, while H3K4me3 may keep the locus primed for stress-dependent induction. This establishes a direct link between chromatin state, BAP1, and CHOP regulation under metabolic stress.

Although BAP1 has been implicated in apoptosis via IP3R3 [10] and ferroptosis via SLC7A11 repression [11], we show that under metabolic stress, hepatocyte death associated with BAP1 loss is characterized by tumor cell-intrinsic upregulation of CHOP and appears to occur independently of inflammatory or necroptotic pathways. Thus, BAP1 emerges as a central regulator of diverse cell death mechanisms, with tissue context dictating the dominant pathway.

While CHOP is classically associated with ER stress-induced apoptosis [40], its role appears to diverge in oncogenic contexts. In BAP1-deficient hepatocytes, CHOP elevation coincides with apoptosis and liver injury. However, when BAP1 loss is combined with YAP activation or PTEN deletion, CHOP remains elevated, yet apoptosis is bypassed and tumorigenesis ensues. In the tumor microenvironment, CHOP contributes to immune evasion by suppressing effector T cell function [56] and promoting the accumulation of myeloid-derived suppressor cells (MDSCs) [57]. More broadly, sustained ER stress has been shown to promote hepatocarcinogenesis through both tumor-intrinsic and immunosuppressive pathways [58,59]. This dual role of CHOP underscores the critical function of upstream regulators such as BAP1 in determining whether ER stress drives cell death or instead fuels tumor-promoting adaptations.

Clinically, these findings carry translational relevance in the context of MASLD and MASH, metabolic conditions characterized by lipid overload, ER stress, and persistent UPR activation [17,18]. Experimental studies suggest that CHOP contributes to the pathological features of MASH, including steatohepatitis and fibrosis [60,61]. As the global incidence of MASLD/MASH-associated HCC continues to rise [30], BAP1-deficient tumors developing in this context may exhibit a heightened dependence on maladaptive UPR signaling. This positions the UPR-CHOP axis as both a biomarker and therapeutic target, unveiling a potentially actionable vulnerability in a defined molecular subset of HCC.

In summary, our work establishes BAP1 as a chromatin regulator that preserves liver homeostasis while functioning as a context-dependent tumor suppressor. By connecting BAP1 loss to lipid imbalance, ER stress activation, and aberrant CHOP induction, we delineate a mechanism through which epigenetic dysregulation rewires stress adaptation in the liver. This work not only expands the functional understanding of BAP1 in liver tumorigenesis but also highlights broader principles by which chromatin modifiers govern cellular responses to metabolic and proteotoxic stress, offering a conceptual framework and potential targets for precision oncology.

## Acknowledgements

We thank all members of the Tschaharganeh lab for constructive discussion and feedback throughout the project. We are grateful to the German Cancer Research Center (DKFZ) Central Animal Laboratory, the Center for Model Systems and Comparative Pathology at the Institute of Pathology Heidelberg, and the DKFZ Genomics Core Facility for excellent technical support. The authors gratefully acknowledges the data storage service SDS@hd supported by the Ministry of Science, Research and the Arts Baden-Württemberg (MWK) and the DFG through grant INST 35/1503-1 FUGG.

This work was funded by ERC Starting Grant “CrispSCNAs” (grant number: 948172) awarded to D.F.T., and by project B05 of the collaborative research grant SFB/TR 209 “Liver Cancer” (grant number: 314905040) through the Deutsche Forschungsgemeinschaft (DFG, German Research Foundation). B.B. is recipient of grant SFB/TRR186-Z4; FOR-5815/1-P3/Z from DFG. D.F.T. is recipient of a TS 293/3-1 from DFG.

## Authors’ contributions

Conceptualization: O.O.G., A.S., D.F.T., M.B.

Methodology: O.O.G., A.S., L.T., A.Sch., D.H., L.Bu., L.W.-L., S.N.

Data acquisition: O.O.G., A.S., L.Bu., L.W.-L., L.B., S.N., D.H.

Analysis and interpretation: A.S., O.O.G., L.P., A.Sch., C.L.

Writing – original draft: O.O.G., A.S., M.B., D.F.T.

Writing – review and editing: all authors

Supervision & funding acquisition: D.F.T., M.B., D.C., M.H., J.Z.-R., B.B

O.G. and A.S. contributed equally. D.F.T. and M.B. are corresponding authors.

## Methods

### Molecular cloning

To perform CRISPR/Cas9-mediated genome editing, sgRNAs were cloned into either the pX330-U6-Chimeric_BB-CBh-hSpCas9 plasmid (Addgene #42230), a vector expressing SpCas9 and chimeric guide RNA, or the lentiCRISPR v2 vector (Addgene #52961), a lentiviral backbone expressing SpCas9 and sgRNA under the U6 promoter, following a previously published protocol [73]. Briefly, BbsI-digested vectors were ligated with annealed and phosphorylated sgRNA oligos, sgGFP (5’-GGGCGAGGAGCTGTTCACCG-3⍰), sgPten (5’-GTTTGTGGTCTGCCAGCTAA-3⍰), sgBap1 (5’-GGGCGAGGAGCTGTTCACCG-3⍰), sgPten (5’-GTTTGTGGTCTGCCAGCTAA-3⍰). Cloned vectors were validated by Sanger sequencing using the human U6 primer.

Effective shRNAs were designed using a published predicition algorithm and cloned into the MSCV-LTR-miR-E-PGK-Puro-IRES-GFP (MLPe) vector as described previously [74, 75]. Two potent murine shRNAs targeting *Bap1* (5⍰-TGCTGTTGACAGTGAGCGACCGAATGAAGGATTTCACCAATAGTGAAGCCACAGATGTATTGGTGAAATCCTTCATTCGGCTGCCTACTGCCTCGG-3⍰ and 5⍰-TGCTGTTGACAGTGAGCGCAAGGAAGAAGTTCAAGATTGATAGTGAAGCCACAGATGTATCAATCTTGAACTTCTTCCTTTTGCCTACTGCCTCGGA-3⍰) and a control shRNA targeting Renilla luciferase (5⍰-TGCTGTTGACAGTGAGCGCAGGAATTATAATGCTTATCTATAGTGAAGCCACAGATGTATAGATAAGCATTATAATTCCTATGCCTACTGCCTCGG-3⍰) were selected based on predicted efficacy. Synthetic 97-mer oligos encoding hairpin structures were PCR-amplified using miRE-XhoI (TACAATACTCGAGAAGGTATATTGCTGTTGACAGTG AGCG) and miRE-EcoRI primers (TTAGATGAATTCTAGCCCCTTGAAGTCCGAGGCAGTAGGCA). The resulting PCR products were digested, purified, and ligated into the EcoRI-HF/XhoI-digested MLPe backbones. All constructs were verified by Sanger sequencing using the miR-E primer.

Target cells were transduced at multiplicity of infection < 0.7 to ensure single-copy integration, followed by a 4-day puromycin selection. Protein lysates were then collected and analyzed by Western blot. The most potent shRNAs, as determined relative to the Renilla-targeting control shRNA, were selected for further experiments.

### Patient samples

Formalin-fixed, paraffin-embedded human liver tumors were obtained from the Institute of Pathology, University of Regensburg (Prof. Dr. Calvisi). Molecular characterization of this cohort, including identification of *BAP1* mutations, has been described previously [62]. Mutational profiling was performed using both the Illumina TruSight Oncology 500 (TSO500) and Qiagen comprehensive cancer gene panels, as detailed in the corresponding publication [62]. All procedures were conducted in accordance with institutional ethical approval as specified therein.

### Analysis of public cancer genomics datasets

Publicly available cancer genomics data were retrieved and analyzed via cBioPortal for Cancer Genomics (https://www.cbioportal.org/). Data from The Cancer Genome Atlas (TCGA) PanCancer Atlas and Liver Hepatocellular Carcinoma (TCGA-LIHC) cohorts were used to assess *BAP1* alterations and their association with *DDIT3* (CHOP) expression. Gene expression comparisons were performed between BAP1 wild-type and mutant tumors.

### Analysis of independent HCC transcriptomic dataset

An independent HCC cohort comprising 406 tumor liver samples was analyzed as part of a previously established project (Pan et al., manuscript in submission). RNA sequencing data were used to evaluate the relationship between *BAP1* status, *DDIT3* expression, and UPR activity. Single-sample gene set enrichment analysis (ssGSEA) was performed using the singscore R package, with the REACTOME_UNFOLDED_PROTEIN_RESPONSE_UPR gene set obtained from the msigdbr database. Statistical significance was determined using the two-tailed Mann-Whitney U test.

### Animal experiments

All animal experiments were conducted at the DKFZ animal facilities. Mice were housed in individually ventilated cages under a 12-h light/dark cycle with free access to food and water.

All animal experiments were conducted in compliance with the regional regulations and with the approval of the regional board (Regierungspräsidium Karlsruhe, Germany), under the permit number: EP-Z160I02, G-178/16-01, G-270/16, G-283/16, A-23/17, G-70/20, and G-81/20.

### Transgenic mice

To model BAP1 loss in the liver, we used a transgenic mouse system incorporating a tetracycline-responsive shRNA construct targeting *Bap1* [21, 22]. The system was generated using embryonic stem cells containing a frt-hygro-pA cassette downstream of the *Col1a1* locus and a reverse tetracycline transactivator (rtTA) inserted into the *Rosa26* locus. Potent shRNAs against *Bap1* or a non-targeting control (Renilla) were inserted into *Col1a1* locus under control of tetracycline-responsive elements (TRE) and a GFP reporter. For liver-specific knockdown, these mice were crossed to B6-Tg(Alb-cre)21Mgn mice expressing Cre recombinase under the albumin promoter. Upon doxycycline administration, rtTA activates shRNA expression in hepatocytes, enabling temporally controlled *Bap1* silencing in the liver. Genotyping was performed using a master mix containing RedTag polymerase, genomic DNA, and gene-specific primers. The *Col1a1* knock-in allele was detected with Col1a1.1 (forward: 5⍰-TTCAGACAGTGACTCTTCTGC-31, reverse: 5⍰-AATCATCCCAGGTGCACAGCATTGCGG-3⍰, internal: 5⍰-CTTTGAGGGCTCATGAACCTCCCAGG-3⍰); the *Rosa26-rtTA* allele with Rosa26.1 (forward 5⍰-AAAGTCGCTCTGAGTTGTTAT-3⍰, reverse: 5⍰-GCGAAGAGTTTGTCCTCAACC-3⍰, internal: 5⍰-CCTCCAATTTTACACCTGTTC-3⍰); and the *Alb-Cre* transgene with Cre (forward: 5⍰-TGCCACGACCAAGTGACAGC-3⍰, reverse: 5⍰-CCAGGTTACGGATATAGTTCATG-3⍰). PCR products were resolved by agarose gel electrophoresis and visualized using a FluorChem M imager (ProteinSimple).

### Hydrodynamic tail vein injection (HTVI)

Female C57BL/6N mice (approximately 8 weeks old, Charles River) were injected with 2⍰ml of sterile saline solution (10% of body weight) containing naked plasmids. For sgRNA- and shRNA-mediated gene disruption 20⍰µg of vectors were used, either pX330-U6-Chimeric_BB-CBh-hSpCas9 (Addgene #42230) carrying sgRNA of interest or pT3-TRE-tRFP-miR-E carrying shRNA of interest. For gene overexpression, 10⍰µg of pT3-EF1a-YAPS127A was injected. The CMV-SB13 plasmid expressing Sleeping Beauty transposase was co-injected at a 1:5 ratio relative to the transposon-based plasmid.

### Subcutaneous cell injection and xenograft measurement

Mice were anaesthetized using isoflurane delivered at a pressure of 1-1.5 bar, with a flow rate of 2-2.5⍰L/min and vaporizer set to 2-2.5 vol%. A total of 0.5⍰× ⍰10^6^ human cells in 100⍰μL PBS were injected subcutaneously into each flank. Tumor growth was monitored by caliper measurements under anaesthesia, and volumes were calculated using the formula: volume = (length × width^2^) / 2.

### Mouse dietary experiments

Mice were fed either a choline-deficient, high-fat diet (CD-HFD; Research Diets D05010402; 45 kcal% fat without added choline), a standard high-fat diet (HFD; Research Diets D12451; 45 kcal% fat), or a doxycycline-supplemented diet (Envigo TD.08541; 6.25% doxycycline hyclate). For combination doxycycline and CD-HFD/HFD treatment, doxycycline-containing water (2 g/L doxycycline and 10 g/L saccharose) was provided. Three treatment regimens were used to control transgene expression: ON (continuous doxycycline), OFF (no doxycycline), and ON-OFF (doxycycline for 2 weeks followed by 4 weeks without). Each experimental group included 5-15 animals unless otherwise specified.

### Serum biochemistry

Blood was collected via submandibular bleeding into serum separation tubes (41.1378.005, Sarstedt). Samples were then centrifuged at 10,000⍰× ⍰g for 10 minutes to isolate serum. Levels of aspartate aminotransferase (AST), alanine aminotransferase (ALT), and total bilirubin (TBIL) were measured using GOT/AST-PIII (9903140), GPT/ALT-PIII (9903150), and TBIL-PIII (9903240) reagents (Fujifilm), using DRI-CHEM NX500i analyzer (Fujifilm, Japan) according to the manufacturer’s instructions.

### Tissue processing

Upon tumor detection by palpation or at the experimental endpoint, mice were sacrificed by cervical dislocation. Livers were inspected under a dissectoscope MZ10 F (Leica) using the Leica Application Suite software to visualize mKate and GFP fluorescence and confirm transgene expression. Tissue samples were snap-frozen in liquid nitrogen for nucleic acid or protein extraction, embedded in OCT compound (Tissue-Tek, Sakura, 4583) for cryosectioning, or fixed in 4% paraformaldehyde for histological analysis. Fixed samples were paraffin-embedded and sectioned at the CMCP (Center for Model Systems and Comparative Pathology, Heidelberg Institute of Pathology, University Clinic Heidelberg) or at the “Chronic Inflammation and Cancer” divison at DKFZ.

### Immunofluorescence

Paraffin-embedded tissue sections were deparaffinized in xylene and rehydrated through graded ethanol solutions (100% ethanol, 96% ethanol and 70% ethanol). After final rehydration step in water, antigen retrieval was performed using a pressure cooker, followed by cooling in running water. Sections were then outlined with a hydrophobic barrier, blocked with 5% BSA in PBS with 0.05% Triton X-100, and incubated with primary antibodies (BAP1, Cell Signaling Technology, 13271S; GFP, Abcam, ab13970; HNF4α, Santa Cruz, sc-6556; KRT19, Abcam, ab52625) overnight at 4°C in a humid chamber. After PBS washes, secondary antibodies (Alexa Fluor 488 donkey anti-chicken, Jackson ImmunoResearch, AB2340375; Alexa Fluor 568 donkey anti-goat, A11058; Alexa Fluor 594 donkey anti-rabbit; Invitrogen, A21207) were applied for 1 hour in the dark. Nuclei were stained by Hoechst (H1398, Invitrogen), followed by mounting medium (ProLong Diamond Antifade Mountant, P36970, Invitrogen), and covered with coverslips. Images were acquired using a Keyence fluorescence microscope, and image processing and analysis were performed using Keyence BZ-X Analyzer.

### Immunohistochemistry

Hematoxylin and eosin (H&E) staining and immunohistochemistry were performed following previously established protocols [76]. TUNEL assays were conducted using the commercially available kits as previously described [77].

Briefly, IHC staining was performed following the same protocol as immunofluorescence (see above) for deparaffinization, rehydration, and antigen retrieval. After blocking, sections were incubated overnight at 4°C with primary antibodies (BAP1, #13271S, CST; CHOP, #5554, CST; CD3, RM-9107-S1 Epredia; F4/80 #123106, BioLegend; RIPK3, ADI-905-242-100BAP1, Enzo). Endogenous peroxidase activity was quenched with 3% hydrogen peroxide (476.1011, CHEMSOLUTE) for 10 minutes. After incubation with HRP-conjugated secondary antibodies (ImmPRESS HRP Goat Anti-Rat IgG Polymer Kit, Peroxidase MP-7404, Vector Laboratories; ImmPRESS HRP Goat Anti-Rabbit IgG Polymer Kit, Peroxidase, MP-7451, Vector Laboratories) for 30 minutes at room temperature, signal was developed using ImmPACT DAB Substrate Kit (SK-4105, Vector Laboratories). Slides were counterstained with hematoxylin (Microscopy Mayer’s hemalum solution, 1.09249.0500, Sigma-Aldrich), dehydrated, and mounted.

Stainings were performed either in-house or at the DKFZ Division of Chronic Inflammation and Cancer. Slides processed at the division were stained on a Leica Bond automated platform using the Leica Bond Polymer Refine Detection Kit (DS9800). The same primary antibodies were applied (except for CD3 where MA1-90582, Invitrogen was used). Blocking steps utilized Leica Bond Primary Antibody Diluent (AR9352). Post-primary and detection steps were performed using reagents included in the Leica Bond system. All slides were mounted with Surgipath Micromount (3801731, Leica), scanned using either the Aperio AT2 system (Leica) for externally processed slides or the NCT Tissue Bank platform for in-house stainings, and analyzed in QuPath.

For human liver tumors, BAP1 and CHOP immunohistochemistry was performed at the Institute of Pathology, University of Regensburg, following previously described procedures [63,64]. BAP1 staining was conducted using an automated procedure (mouse monoclonal anti-BAP1, clone C-4, sc-28383, Santa Cruz Biotechnology), while CHOP staining followed a manual protocol (rabbit polyclonal anti-CHOP, 15204-1-AP, Proteintech) with anti-mouse secondary antibody (ZUC 032-100, Zytomed). Signals were developed using the ImmPACT NovaRed Substrate/Chromogen Kit (SK-4805, Vector Laboratories), slides were mounted with Leica mounting medium and scanned using a 3D Histech P1000 scanner.

### Extraction and quantification of nucleic acids and proteins

Plasmid DNA was isolated using the QIAGEN Plasmid Plus Midi Kit (Qiagen, 12945) or the QIAprep Spin Miniprep Kit (Qiagen, 27106), and DNA fragments were purified using the QIAquick Gel Extraction Kit (Qiagen, 28706X4) or the QIAquick PCR Purification Kit (Qiagen, 28106), following the manufacturer’s instructions. Genomic DNA was extracted using the Puregene Core Kit A (Qiagen, 158445). Total RNA was extracted using the RNeasy Mini Kit (Qiagen, 74106).

Nucleic acid concentrations were measured using a NanoDrop spectrophotometer (Thermo Fisher). Protein concentrations were measured using the Bradford Protein Assay Kit (Bio-Rad, 5000006), according to the manufacturers’ protocols.

### Native chromatin immunoprecipitation

Cells were lysed in ice-cold buffer (10⍰mM Tris-HCl pH 8.0, 10⍰mM NaCl, 3⍰mM MgCl_2_), followed by sequential extraction using buffers containing 0.1% NP-40 and 5⍰mM N-ethylmaleimide (NEM). Nuclei were digested with 150-200⍰U MNase (Thermo Scientific, EN0181) for 5 minutes at 37⍰°C, then quenched with 1⍰M EDTA. Supernatants were collected and pooled. Chromatin was either snap-frozen or processed immediately.

For ChIP, 100⍰µl chromatin was diluted 1:10 in buffer (70⍰mM NaCl, 10⍰mM Tris-HCl pH 7.5, 2⍰mM MgCl_2_, 2⍰mM EDTA, 0.1% Triton X-100, 10⍰mM NEM, 1× Thermo Scientific Halt™ Protease Inhibitor Cocktail), centrifuged, and incubated overnight at 4⍰°C with antibodies: ubiquityl-histone H2A (Cell Signaling Technology, #8240) or tri-methyl-histone H3 (Cell Signaling Technology, #9751), or Normal Rabbit IgG (Cell Signaling Technology, #2729) as a negative control. Input was taken prior to antibody addition.

Blocked protein A beads (IPA300, RepliGen) were added and rotated for 1 hour at 4⍰°C. Complexes were washed 4× with wash buffer (20⍰mM Tris-HCl pH 7.5, 2⍰mM EDTA, 125⍰mM NaCl, 0.1% Triton X-100) and once with TE. DNA was eluted in 1% SDS/0.1⍰M NaHCO_3_, purified (Zymo ChIP DNA Clean & Concentrator), and eluted in EB.

### Crosslinked chromatin immunoprecipitation

Cells were fixed with 1% PFA for 10 minutes at room temperature and quenched with 125⍰mM glycine. After PBS washes, cells were lysed in buffer (50⍰mM HEPES-KOH pH 7.5, 140⍰mM NaCl, 1⍰mM EDTA, 1% Triton X-100, 0.1% sodium deoxycholate, 0.1% SDS, 1⍰mM PMSF, 1× protease inhibitor cocktail). Lysates were sonicated using a Bioruptor (Diagenode) to achieve DNA fragment sizes between 200-500 bp. Debris was cleared by centrifugation, and the supernatant was pre-cleared with Protein G magnetic beads for 1 hour at 4⍰°C. An input sample was collected prior to immunoprecipitation.

Chromatin was incubated overnight at 4°C with antibodies: FLAG® M2 (F1804,MilliporeSigma) or Normal Rabbit IgM (#77591, Cell Signaling Technology). Protein G magnetic beads were added and incubated for 2 hours at 4⍰°C with rotation. Complexes were first washed 2× with first wash buffer (50⍰mM Tris-HCl pH 8.0, 150⍰mM NaCl, 5⍰mM EDTA, 1% NP-40, 0.5% sodium deoxycholate, 0.1% SDS), followed by 4× with second wash buffer (100⍰mM Tris-HCl pH 8.5, 500 mM LiCl, 1% NP-40, 1% sodium deoxycholate), 2× with first wash buffer, and then finally 2× with TE buffer (10⍰mM Tris-HCl pH 8.0, 1⍰mM EDTA). DNA was eluted (1% SDS, 0.1⍰M NaHCO_3_), reverse crosslinked overnight at 65⍰°C, purified (QIAquick PCR purification kit, Qiagen), and eluted in EB.

### Chromatin immunoprecipitation qPCR (ChIP-qPCR)

For qPCR, ChIP DNA was diluted to represent 2% of the input. Reactions were performed in triplicate using SYBR Green and gene-specific primers for CHOP (forward: 5⍰-TGCCACTTTCTGATTGGTAGGTT-3⍰; reverse: 5⍰-TGCCACCCGCTCATCTTT-3⍰) on a QuantStudio 3 Real-Time PCR System (Thermo Fisher). Enrichment was calculated relative to input DNA.

### Real-time quantitative PCR (RT-qPCR)

Upon total RNA extraction, cDNA was synthesized with the TaqMan Reverse Transcription Kit (N8080234, Applied Biosystems). Diluted cDNA was used for qPCR with SYBR Green Master Mix (4367659, Thermo Fisher Scientific) and gene-specific primers: BAP1 (forward: 5⍰-CCCCGCGGGAAGATGAATAA-3⍰, reverse: 5⍰-ACCCCCTTGACACCGAAATC-3⍰, DDIT3 (forward: 5⍰-CCTTTCTCCTTCGGGACACT-31, reverse: 5⍰-TTGATTCTTCCTCTTCATTTCCAGG-31), ACTB (forward: 5⍰-CATGTACGTTGCTATCCAGGC-3⍰, reverse: 5⍰-CTCCTTAATGTCACGCACGAT-3⍰). Reactions were run in triplicates on a QuantStudio 3 (Thermo Fisher), an expression was normalized to housekeeping gene expression using the ΔΔCt method.

### Immunoblotting

Cells were lysed, followed by protein concentration estimation. Equal amounts of denatured protein were separated by SDS-PAGE and transferred to PVDF membrane (120V, 90 minutes). Membranes were blocked in 5% milk in TBS-T and incubated with primary antibodies (BAP1, Cell Signaling Technology, 13271S; CHOP, Cell Signaling Technology, 5554S; GFP, Cell Signaling Technology, 2956; Actin-HRP, Sigma-Aldrich, A3854) overnight at 4°C. Following TBS-T washes, secondary antibodies were applied for 1 hour. Signal was detected using an ECL solution (Clarity Western ECL Substrate, 170-5060, Bio-Rad) and visualized with the FluorChem M (ProteinSimple) imager.

### Cell culture

All cells were cultured at 37°C with a 5% CO_2_ incubator. Complete DMEM (Dulbecco’s Modified Eagle Medium, D6429, Sigma-Aldrich) supplemented with 10% FBS (fetal bovine serum) and 1% penicillin-streptomycin was used as the growth medium. Mouse cell lines were cultured on collagen-coated plates (PureCol, 0.05 mg/mL, 5005, CellSystems).

The human cell lines used for experiments included HEK293T, Huh7, Hep3B, and HepG2. For ER stress induction, HCC cell lines (Huh7, Hep3B, HepG2) were treated with tunicamycin (1:1000 dilution, J67401.XF, Thermo Scientific Chemicals) for 24 hours.

HEK293T cells were transfected with 50⍰µg of expression vector using PEI (1⍰mg/mL) in 15⍰cm dishes. After 24 hours, cells were treated with tunicamycin (1:1000) for 4 hours and harvested.

### Primary cell line derivation

Liver tumors were resected using sterile instruments, finely minced with a scalpel, and incubated in digestion medium (prepared in serum-free DMEM) containing 4⍰mg/mL Collagenase IV (Sigma-Aldrich, 11088866001) and Dispase (Sigma-Aldrich, 4942078001) at 37⍰°C. After enzymatic digestion, cells were centrifuged at 300 g for 5⍰minutes, and the supernatant was replaced with complete DMEM. Once cell lines were established, routine mycoplasma testing was performed.

### Virus production and transduction

Lentiviral particles were produced in HEK293T cells by plasmid transfection using PEI (8 µg psPAX2, 2.5 µg pMD.2G, 10 µg vector). Supernatants were collected 48 hours post-transfection, filtered (0.45 µm cellulose acetate membrane), and used to transduce target cells in the presence of 4 µg/ml polybrene (Sigma-Aldrich, 107689). Selection with puromycin (Thermo Fisher Scientific, BP2956-100, 2 µg/ml) began 3 days post-transduction.

### Transcriptomics and lipidomics analyses

RNA sequencing (RNA-seq) was conducted at the DKFZ sequencing core facility. Following alignment of the RNA-seq counts, further analysis was executed using the HUSAR platform at DKFZ. The HUSAR program HTSeq count was employed for genome-wide transcript counting, resulting in FPKM (fragments per kilobase of transcripts per million) values. Differentially expressed genes were identified using DEseq2 within the COMPARNA pipeline. Each condition was analyzed using⍰3 biological replicates. Data were used for downstream pathway analyses (Ingenuity Pathway Analysis (IPA) and Gene Set Enrichment Analysis (GSEA)).

For lipidomics, approximately 50 ng of snap frozen liver tissue was transferred to MK28-R/2ml for subsequent lysis. Samples were processed at the Heidelberg University Biochemistry Center (BZH) at prof. Britta Brueger’s lab. Briefly, cells were subjected to lipid extractions using an acidic liquid-liquid extraction method [65]. On average, 42.5 µg liver (wet weight) was subjected to extractions. Quantification was achieved by adding internal lipid standards for each lipid class. Lipid standards were added prior to extractions, using a master mix consisting of 100 pmol deuterated cholesterol (D_6_-cholesterol, Cambridge Isotope Laboratory), 25 pmol diacylglycerol (DG, 17:0/17:0, Larodan), 25 pmol cholesteryl ester (CE, 9:0, 19:0, 24:1, Sigma), and 24 pmol triacylglycerol (TG, mix of LM-6000 and D_5_-17:0,17:1,17:1, Avanti Research). The final chloroform phase was evaporated under a gentle stream of nitrogen at 37°C. Samples were either directly subjected to mass spectrometric analysis, or were stored at −20°C prior to analysis, which was typically done within 1-2 days after extraction. Lipid extracts were resuspended in 10 mM ammonium acetate in 60 µl methanol. 2 µl aliquots of the resuspended lipid extracts were diluted 1:10 in 10 mM ammonium acetate in methanol in 96-well plates (Eppendorf twin tec 96) prior to measurement. For cholesterol determinations, the remaining lipid extract was again evaporated and subjected to acetylation as described in [66]. Samples were analyzed on an QTRAP 6500+ mass spectrometer (Sciex) with chip-based (HD-D ESI Chip, Advion Interchim Scientific) electrospray infusion and ionization via a Triversa Nanomate (Advion Interchim Scientific) as described. MS settings and scan procedures are listed in Supplementary Table 1. Data evaluation was done using LipidView 1.3 (Sciex) and an in-house-developed software (ShinyLipids). The amount for endogenous molecular lipid species was calculated based on the intensities of the internal standards.

### Patient and public involvement

The public was not involved in this research project.

### Illustrations

Figures, including the graphical abstract (BioRender.com/exgoujf), Figure 1A (BioRender.com/sz8w2to), and Figure 2D (BioRender.com/2k8jqk0), were created using BioRender.com under an academic license.

### Statistical analysis

Statistical analyses were carried out using GraphPad Prism 8. Error bars indicate standard deviation. The specific statistical analysis used for each figure is indicated in the respective figure caption. Significance was denoted as follows: * for p < 0.05, ** for p < 0.01, *** for p < 0.001, **** for p < 0.0001; non-significant results are unmarked.

---

**Supplementary Figure 1.**
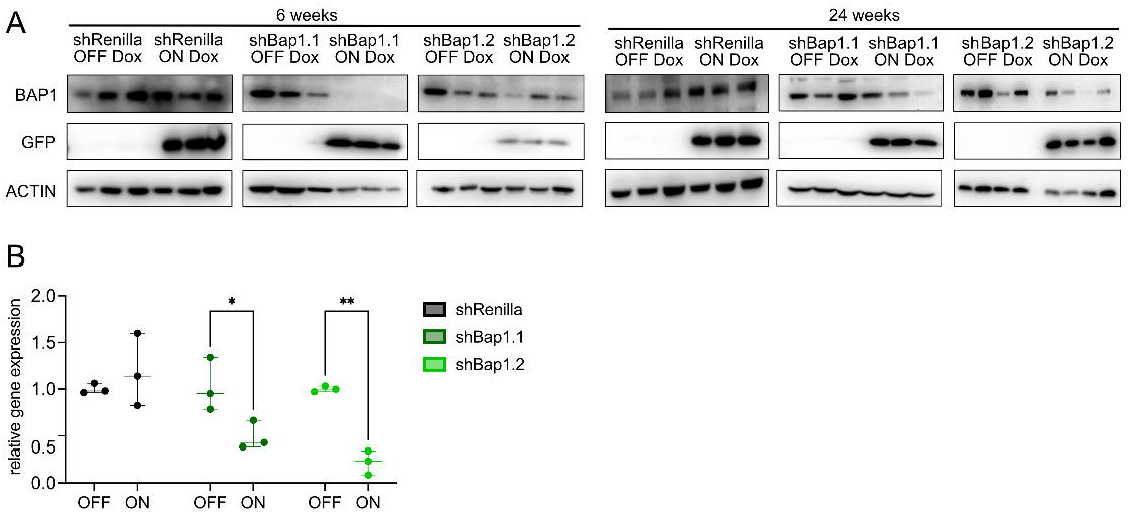
Efficient long-term *Bap1* knockdown *in vivo*. (A) Western blot analysis of whole liver lysates showing diminished BAP1 protein levels in shBap1.1 and shBap1.2 mice after 6 and 24 weeks of Dox treatment. GFP expression correlates with shRNA induction. (B) *Bap1* transcript expression quantification at 6 weeks confirming reduced *Bap1* gene expression in both shBap1 strains under Dox. N = 3, two-way ANOVA.

**Supplementary Figure 2.**
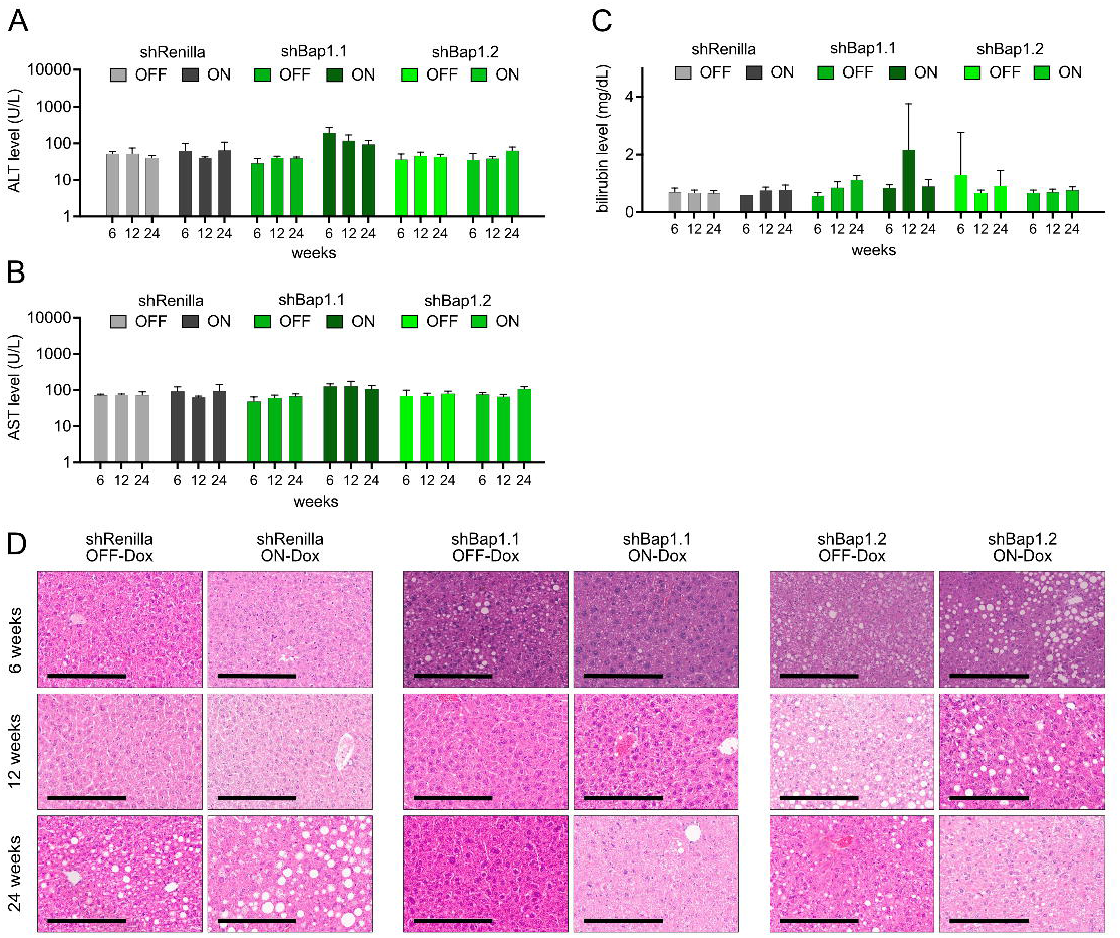
Long-term *Bap1* knockdown is well tolerated under basal conditions. (A-C) Quantification of serum liver function markers in Dox-treated shBap1.1, shBap1.2, and shRenilla mice at 6, 12 and 24 weeks: (A) ALT, (B) AST, and (C) bilirubin levels remained within normal range across all groups. N ≥ 3, two-way ANOVA with Tukey’s test. (D) Representative liver histology (H&E) showing preserved architecture with occasional hepatocellular vacuolization but no evidence of neoplastic transformation, scale bar: 200 µm.

**Supplementary Figure 3.**
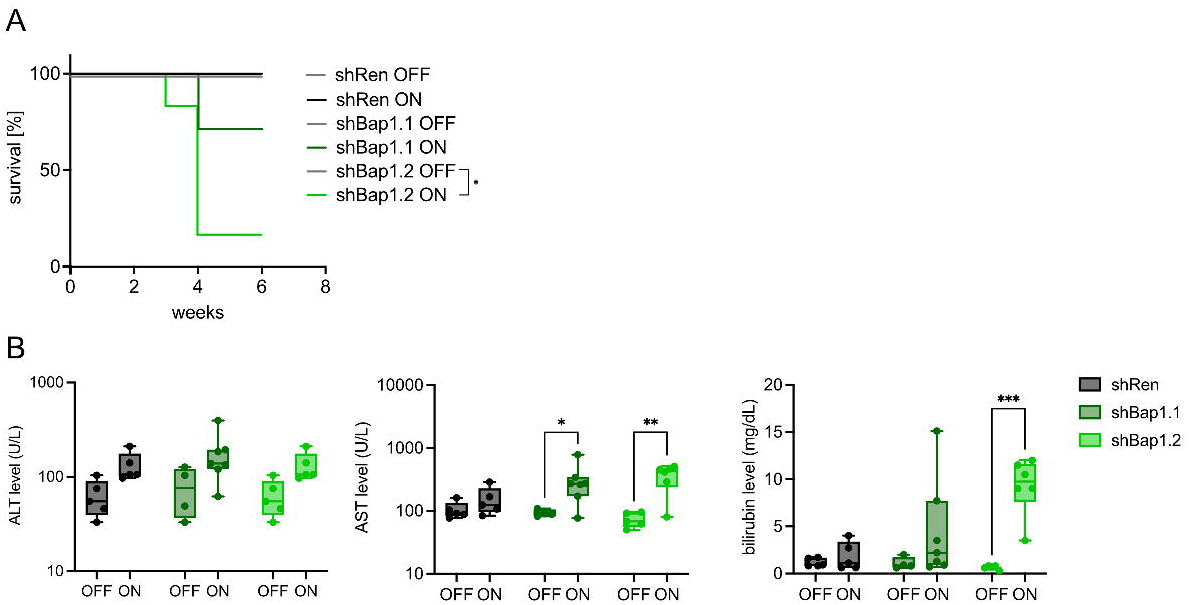
Liver failure in BAP1-deficient HFD-fed mice. (A) Kaplan-Meier survival curves of shBap1.1, shBap1.2, and shRenilla mice treated with HFD under continuous Dox (ON) or no Dox (OFF). N ≥ 4, Log-rank (Mantel-Cox) test. (D) Serum levels of ALT, AST, and bilirubin measured at endpoint show elevated liver injury markers in BAP1-deficient mice. N ≥ 4, two-way ANOVA with Šídák’s test.

**Supplementary Figure 4.**
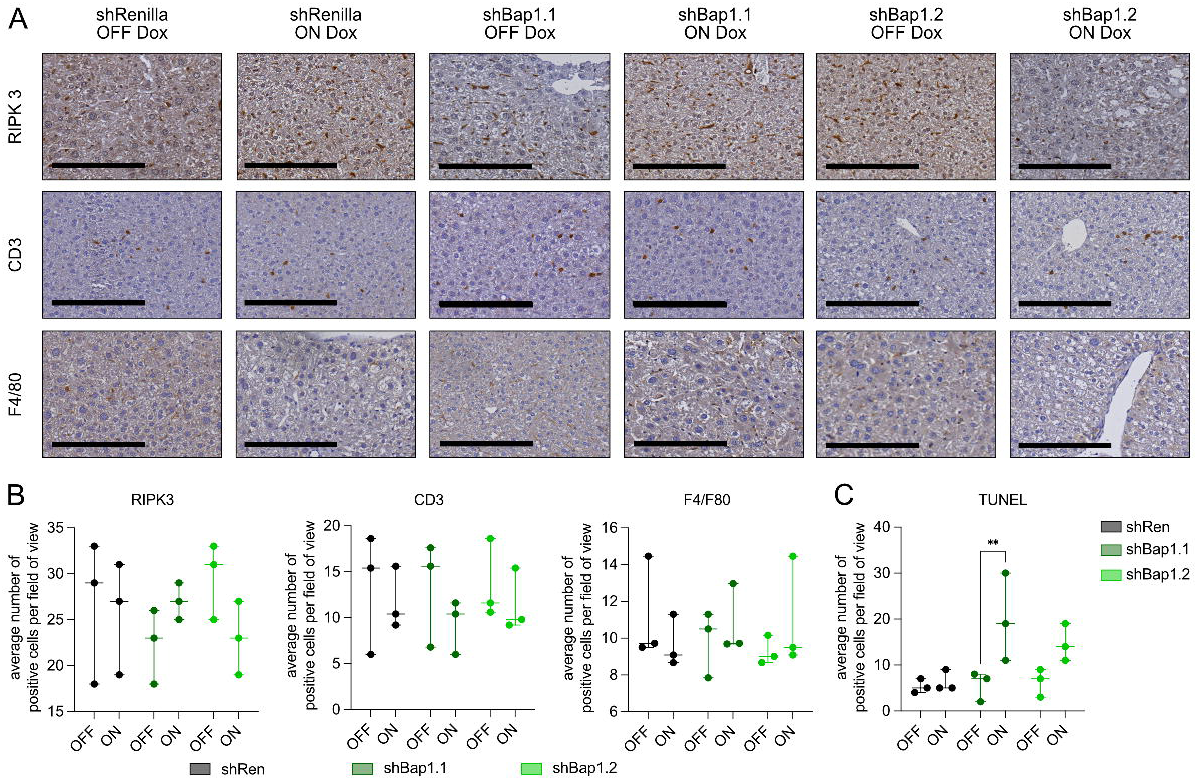
BAP1 loss does not trigger immune infiltration or necroptosis under metabolic stress. (A&B) Representative immunohistochemistry images (A) and associated quantifications (B) of RIPK3, CD3, and F4/80 staining reveal no major differences across groups, indicating no significant immune recruitment or necroptotic signaling; scale bar: 200⍰µm. (C) Quantification of TUNEL^+^ cells shows increased apoptosis in Dox-treated shBap1 livers. N = 3, two-way ANOVA with Šídák’s test.

**Supplementary Figure 5.**
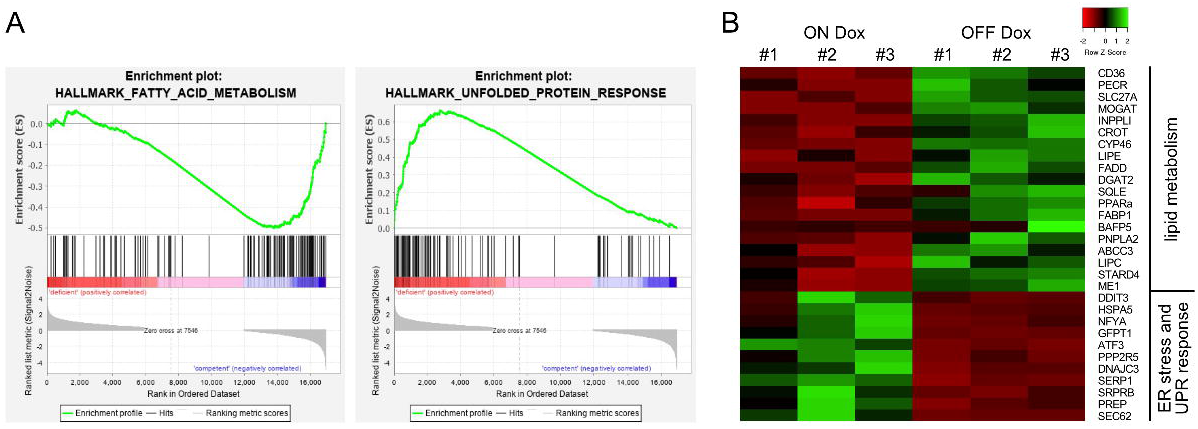
BAP1 loss represses fatty acid metabolism and activates the UPR. (A) Gene Set Enrichment Analysis (GSEA) of liver RNA-seq comparing ON Dox (shBap1) versus OFF Dox shows negative enrichment of fatty acid metabolism and positive enrichment of unfolded protein response. (B) Heatmap of representative lipid metabolism and ER stress/UPR genes across biological replicates; values shown as row z-scores (scale shown). N = 3 per group.

